# Staphyloxanthin production by *Staphylococcus aureus* promotes resistance to oxidative stress to delay diabetic wound healing

**DOI:** 10.1101/2022.09.09.507290

**Authors:** Amelia R. McCready-Vangi, Amy E. Campbell, Aayushi Uberoi, Victoria M. Lovins, Ellen K. White, Alexis R. Morgenstern, Sue E. Gardner, Elizabeth A. Grice

**Author notes:** Contributed equally (Amelia R. McCready-Vangi, Amy E. Campbell).

## Abstract

Diabetic foot ulcers (DFU) are a serious complication of diabetes mellitus that burden patients and health care systems. *Staphylococcus aureus* is prevalent and abundant in the DFU microbiome, and strain-level differences in *S. aureus* may drive clinical outcomes. To identify mechanisms underlying strain-specific outcomes in DFU with *S. aureus*, we performed high-throughput phenotyping screens on a collection of 221 *S. aureus* cultured isolates from clinically uninfected DFU. Of the 4 phenotypes examined (*in vitro* biofilm formation and production of staphylokinase, staphyloxanthin, and siderophores), we discovered that isolates from non-healing wounds produced more staphyloxanthin, a carotenoid cell membrane pigment. In a murine diabetic wound healing model, staphyloxanthin-producing isolates delayed wound closure significantly compared to staphyloxanthin-deficient isolates. Staphyloxanthin promoted resistance to oxidative stress *in vitro* and enhanced bacterial survival in human neutrophils. Comparative genomic and transcriptomic analysis of genetically similar clinical isolates with disparate staphyloxanthin phenotypes revealed a mutation in the Sigma B regulatory pathway that resulted in marked differences in stress response gene expression. Our findings suggest that staphyloxanthin production delays wound healing by protecting *S. aureus* from neutrophil-mediated oxidative stress, and may provide a target for therapeutic intervention in *S. aureus*-positive wounds.

## INTRODUCTION

Non-healing wounds are a common, costly, and debilitating complication of diabetes. Lifetime incidence of a diabetic foot ulcer (DFU) is estimated at 19-34% in those with diabetes (Armstrong et al., 2017). Treatment of lower extremity complications are responsible for ~1/3 of total diabetic care costs, and standard treatment approaches include surgical debridement, offloading, and often local or systemic antimicrobials (Everett and Mathioudakis, 2018; Hartemann-Heurtier and Senneville, 2008). However, prognosis for patients with DFU remains poor, and DFU account for two-thirds of all non-traumatic amputations performed in the United States (Hoffstad et al., 2015; Martins-Mendes et al., 2014). A DFU strongly increases risk of death in those with diabetes, with five-year mortality rates of 40-50% (Armstrong et al., 2007; Moulik et al., 2003). Despite the global healthcare burden posed by DFU complications, clinical management strategies have not evolved to meet the escalating need.

Many DFU complications owe to infections by contaminating bacteria which colonize the wound bed. Of these microbes, which may originate from the adjacent skin microbiome or surrounding environment, the Gram-positive pathogen *Staphylococcus aureus* is the predominant bacterial species found in DFU (Dunyach-Remy et al., 2016; Gardiner et al., 2017). Highly prevalent across many types of cutaneous wounds, *S. aureus* is an important contextual pathogen that causes local and systemic infections that are difficult to treat (Citron et al., 2007; Sloan et al., 2019; Tong et al., 2015; Wolcott et al., 2016). Treatment for *S. aureus* is made more complicated by its evolving resistance to currently available antibiotics including methicillin and vancomycin (Dunyach-Remy et al., 2016). Still, *S. aureus* can asymptomatically colonize the skin and nares, and its presence in a DFU does not necessitate infection (Gardner et al., 2014; Olaniyi et al., 2017).

Rather, prior studies indicate that *S. aureus* effects on DFU healing are strain dependent, and that uninfected and infected DFU can be differentiated by virulence traits of colonizing *S. aureus* strains which enable them to survive in the cutaneous wound environment and evade host immune responses (Sotto et al., 2008, 2012). Using shotgun metagenomics, we previously showed that specific strains of *S. aureus* populate healing vs non-healing DFU and that *S. aureus* exerts strain-specific effects on wound pathogenesis in a diabetic murine model (Kalan et al., 2019). In the same study, metagenomic profiles from nonhealing DFU were enriched for *S. aureus* virulence- and pathogenicity-associated genes, including biofilm formation, as compared to healing DFU. These and prior findings suggest that strain-level differences in *S. aureus* virulence factors contribute to delayed healing and other complications in DFU.

*S. aureus* possesses an arsenal of virulence traits which vary between genetic lineages and may therefore help certain strains to infect the wound bed and impede healing. Though these virulence traits include secreted toxins which directly damage host tissues, they also include mechanisms by which *S. aureus* survives attacks from host cells and proliferates despite host-imposed limitations to bacterial survival. Biofilms, for example, entrench *S. aureus* in a resilient extracellular matrix, protecting it from antimicrobials and clearing by the host immune system (Roy et al., 2014; Schierle et al., 2009). Siderophores, which competitively bind environmental iron for uptake by *S. aureus*, enable it to overcome nutrient limitations inflicted by host iron sequestration (Dale et al., 2004; Harrison et al., 2008). To evade innate immune detection and clearance, *S. aureus* produces factors such as the anti-opsonic protein staphylokinase. Staphylokinase protects *S. aureus* from host immune responses by converting host plasminogen to plasmin, which breaks down fibrin clots and clears opsonins such as immunoglobulin G (IgG) and C3b from the wound bed (Rooijakkers et al., 2005). In the highly oxidative diabetic wound environment (Cano Sanchez et al., 2018; Dworzański et al., 2020), *S. aureus* can suffer substantial damage to its proteins and genetic material. To prevent and mitigate this damage, *S. aureus* deploys stress resistance factors such as the carotenoid pigment staphyloxanthin, which neutralizes reactive oxygen species such as those produced by host neutrophils (Clauditz et al., 2006; Liu et al., 2005). Such virulence factors are likely essential to *S. aureus* survival and proliferation in the diabetic would environment. However, it remains unclear which protective phenotypes most strongly determine DFU outcomes, and how these phenotypes enable certain strains of *S. aureus* to cause tissue damage and delay healing in a diabetic wound context.

To address this gap, we employed a collection of 221 *S. aureus* clinical isolates cultured from a longitudinal prospective study of wound bioburden in 100 neuropathic DFU (Gardner et al., 2014). We characterized virulence in the resulting *S. aureus* isolates using high-throughput screens for traits which confer *S. aureus* resistance to host immune attacks. We identified an association between DFU healing outcome and *in vitro* production of the antioxidant pigment staphyloxanthin. Leveraging clinical isolates with variable staphyloxanthin phenotypes and transposon mutants for staphyloxanthin production, we found that staphyloxanthin delayed wound healing in a diabetic murine model. Staphyloxanthin-producing strains enhanced neutrophil recruitment, were more resistant to oxidative stress, and survived intracellularly in neutrophils. Using whole genome sequencing and comparative genomics of high and low staphyloxanthin-producing isolates, we identified a mutation which attenuated staphyloxanthin production via a regulator of Sigma B, an alternative sigma factor involved in *S. aureus* stress response. Transcriptomic comparisons revealed that in response to peroxide stress, a clinical isolate with the mutation exhibited greater upregulation of oxidative stress response genes and protein damage repair genes compared to a high-staphyloxanthin clinical isolate lacking the mutation.

## RESULTS

### Phenotypes of *S. aureus* in a culture collection from human diabetic foot ulcers

We used a collection of *S. aureus* isolates from a previously described longitudinal prospective cohort study of DFU bioburden and association with clinical outcomes (Gardner et al., 2014). Enrolled subjects presented with plantar, neuropathic DFU lacking signs of clinical infection and underwent sharp debridement followed by offloading. DFUs were monitored bi-weekly for 26 weeks or until healed or amputated, with wound swab specimens collected at each visit (**Figure 1A**). Swabs were plated under a range of media conditions, including *S. aureus-*selective chromogenic detection plates. Sixty DFU were positive for *S. aureus* at some time point during the 26-week study, yielding 221 total *S. aureus* isolates. Of the 60 DFU that were *S. aureus* culture-positive, 43 healed by the end of the study (72%) (**Figure 1A**). Roughly similar numbers of *S. aureus* isolates were obtained from DFU that healed (n=108) versus DFU that did not heal by the end of the study (n=113).

**Figure 1:**
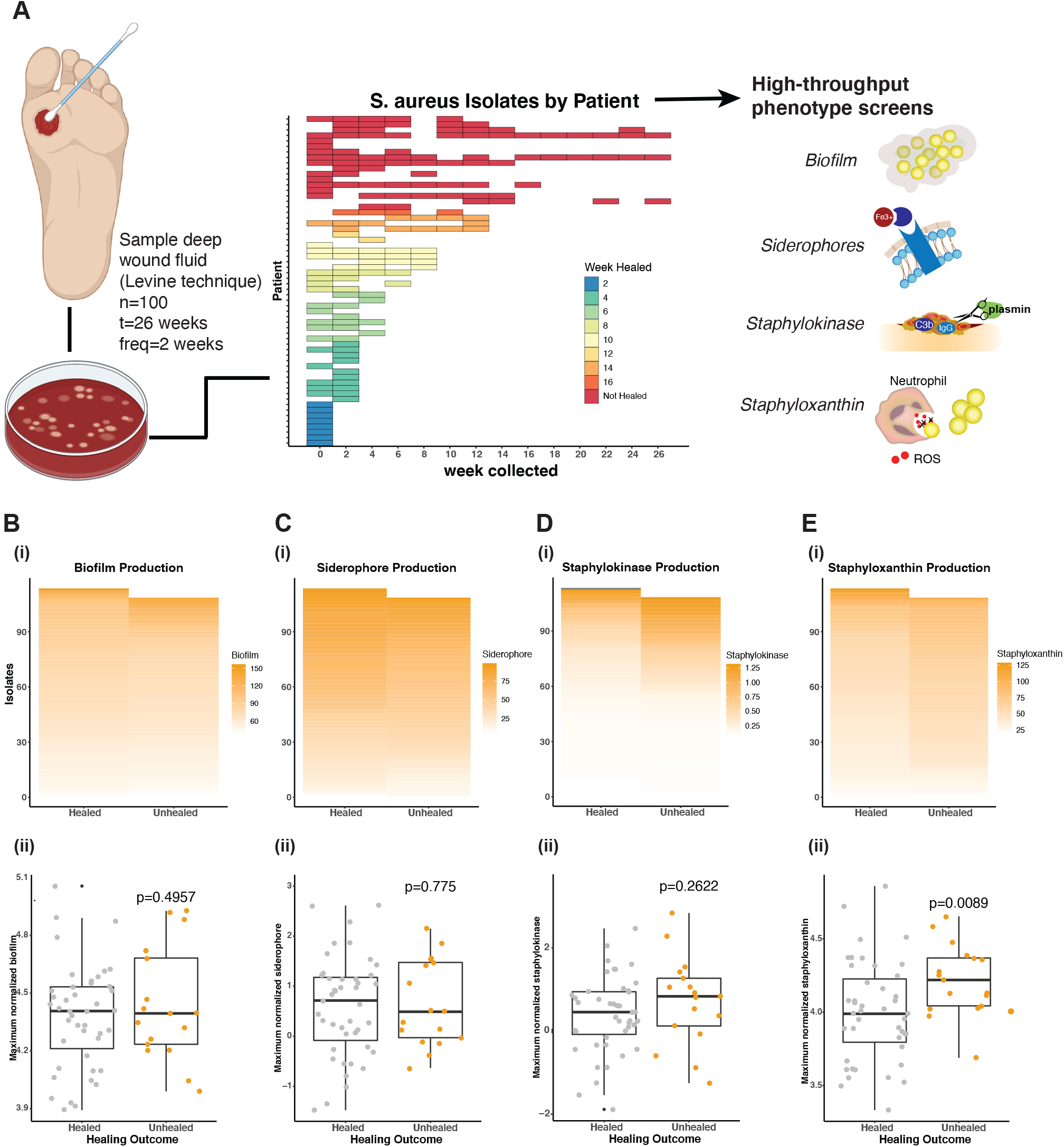
*Staphylococcus aureus* virulence phenotypes are associated with diabetic foot ulcer outcomes. **(A)** The Levine technique was used to collect swab specimens from diabetic foot ulcers (DFU; n=100) and cultured on selective agar to identify *S. aureus*. Of the 100 DFU, 60 were culture-positive for *S. aureus*. Variable virulence phenotypes were measured in all resulting isolates (n=221) by high throughput screens, where measures-by-isolate represent the means of 3 biological replicates by isolate averaged from 3 technical replicates each. **(B-E, i panels)** Distribution of phenotypes by isolate for all *S. aureus* isolates from healed (left) and unhealed (right) DFU by the end of the 26-week study. **(B-i)** Biofilm production (% of *S. aureus* 113 mean), **(C-i)** siderophore production (% of *S. aureus* 502A mean), **(D-i)** staphylokinase production (ug)**, (E-i)** staphyloxanthin production (% of *S. aureus* 502A mean). **(B-E, ii panels)** Distribution of normalized virulence factor production (y-axis) by a single representative isolate per DFU, as chosen by the isolate with the highest virulence factor production for each DFU, healed (left) and unhealed (right) by study end. P-values are reported from t-tests on representative isolate measure ~ healing outcome for **(B-ii)** log-normalized biofilm production, **(C-ii)** order quantile-normalized siderophore production, **(D-ii)** order quantile-normalized staphylokinase production, and (**E-ii)** log-normalized staphyloxanthin production. See also Table S1, S2 and Figure S1.

To characterize variation in the 221 *S. aureus* isolates in this collection, we performed high-throughput screens for four stress- and immune-responsive phenotypes we hypothesized to segregate with wound outcomes: biofilm formation (**Figure 1B**), siderophore production (**Figure 1C**), staphylokinase activity (**Figure 1D**), and staphyloxanthin production (**Figure 1E**). The distribution of phenotypes by isolate for all *S. aureus* isolates from healed (left) and unhealed (right) DFU by the end of the 26 week study are depicted in **Figure 1B-E, i panels** (see also **Table S1**). Since some isolates came from the same DFU, and/or timepoint, we de-duplicated the dataset for statistical analysis by selecting the highest-virulence isolate for each virulence phenotype in each DFU.

We assessed *in vitro* biofilm formation using an adapted crystal violet staining technique (O’Toole, 2011). Although *S. aureus* isolates varied in ability to form biofilm, isolates obtained from non-healing DFU did not produce significantly more biofilm *in vitro* than isolates from DFU that healed by the end of the 26-week study (p=0.4957; **Figure 1Bii**). Siderophore production, quantified by measuring iron depletion in the presence of *S. aureus* supernatants (Schwyn and Neilands, 1987), was also variable but did not differ between healed and unhealed DFU (p=0.775; **Figure 1Cii**).

We quantified staphylokinase production in all isolates using a plasmin-specific chromogenic substrate that is activated when staphylokinase cleaves exogenously added human glu-plasminogen into plasmin (Kwiecinski et al., 2013). We observed that there was no significant difference in staphylokinase production when comparing isolates from healed and unhealed DFU (p=0.2622; **Figure 1Dii**).

The carotenoid pigment staphyloxanthin was extracted from liquid cultures via methanol and quantified colorimetrically. Isolates from non-healing DFU produced significantly more staphyloxanthin compared to isolates from DFU that healed (p=0.0089; **Figure 1Eii**). Similar results were obtained when comparing isolates from DFU that did or did not heal by 12 weeks, a timepoint which is commonly used to distinguish chronic nonhealing DFU (p=0.00418) (**Figure S1**).

In addition to comparing each phenotype in deduplicated isolates from healing and nonhealing patients using t-tests (**Figures 1B-E, ii panels**), we fit a multilevel linear mixed model (LMM) for each phenotype of the form *Phenotype ~ (1*|*patient) + HealingOutcome*, where *Phenotype* is the min-max normalized, mean phenotype measure for an isolate across biological and technical replicates, *patient* is patient ID, and *HealingOutcome* represents healed or nonhealed status at the end of the study (Bates et al., 2015). Of the four phenotype models, staphyloxanthin displayed the strongest relationship to the healing outcome factor of the four phenotypes in a well-fitting model (**Table S2**). Collectively, these data demonstrate an association between increased staphyloxanthin production by *S. aureus* and poor clinical outcome of DFU.

### Staphyloxanthin promotes *S. aureus* survival under oxidative conditions

To further examine the association between staphyloxanthin and delayed healing, we looked at the distribution of staphyloxanthin production among isolates from each individual DFU (**Figure 2A**). One non-healing DFU in particular, DFU141, presented with a unique staphyloxanthin profile: of the 14 *S. aureus* isolates obtained from DFU141 over the course of 16 weeks, 11 produced distinctly high and 3 produced distinctly low levels of staphyloxanthin (**Figure 2B**). This observation among DFU141 *S. aureus* isolates motivated a case study of dichotomous staphyloxanthin production in a subset of our clinical isolates. We selected representative isolates with highest and lowest staphyloxanthin production from this DFU for in-depth analysis: SA925 and SA1088, respectively (**Figure 2B**).

**Figure 2:**
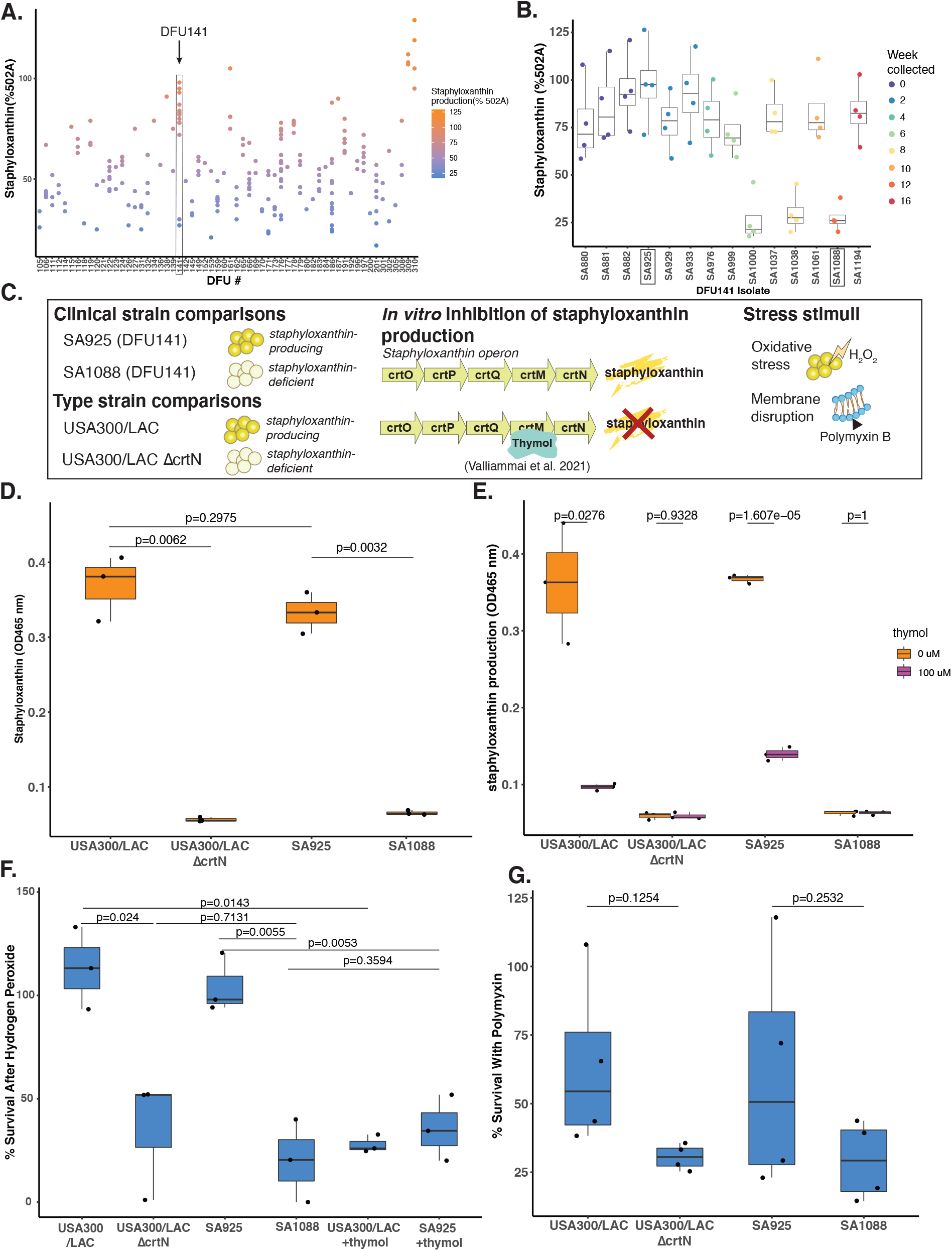
Staphyloxanthin promotes *S. aureus* growth and survival under oxidative stress. **(A)** Distribution of staphyloxanthin production (y-axis) from all individual isolates cultured from each DFU (x-axis), with DFU141 isolates outlined. **(B)** Staphyloxanthin production (% 502A) of each isolate cultured from DFU141 for each of three biological replicates (x-axis). **(C)** Schematic of comparison strains and conditions in (D-E). **(D)** Staphyloxanthin production (mean OD465 across technically replicates) for 3 biological replicates of each comparator strain. **(E)** Staphyloxanthin production in each strain with (right) and without (left) the addition of 100uM thymol. **(F)** % Survival of USA300/LAC, LACΔcrtN, SA925, and SA1088 strains treated with 3% H_2_O_2_. **(G)** % Survival following Polymyxin B treatment. **(A-G)** Each dot represents the mean of 3 technical replicates for 1 biological replicate. Independent t-tests were performed to compare pairs of isolates. See also Figure S2.

To isolate the effects of staphyloxanthin alone, we obtained *S. aureus* isolate USA300 LAC and a transposon mutant of the *crtN* gene in parent strain (USA300 LAC *ΔcrtN;* **Figure 2C**) (Fey et al., 2013). The *crtN* gene encodes the dehydrosqualene desaturase enzyme, a critical enzyme in staphyloxanthin biosynthesis and part of the *crt* operon which encodes for staphyloxanthin production (Pelz et al., 2005). A *crtN* transposon mutant produces virtually no staphyloxanthin, and significantly less than USA300 LAC (p=0.0062; **Figure 2D**). Similarly, in the clinical isolate comparison, SA925 produced significantly more staphyloxanthin than SA1088 (p=0.0032; **Figure 2D**). To inhibit staphyloxanthin production in clinical isolates, we used thymol (2-Isopropyl-5-methylphenol), a monoterpene alcohol isolated from the plant *Thymus vulgaris*. Thymol selectively inhibits staphyloxanthin production in *S. aureus* via binding to CrtM (**Figure 2C**), the first enzyme in the staphyloxanthin biosynthesis pathway (Valliammai et al., 2021). The addition of thymol (100μM) robustly abrogated staphyloxanthin production in both high staphyloxanthin producing isolates, USA300 LAC and SA925 (**Figure 2E**).

Staphyloxanthin promotes *S. aureus* survival by providing protection from oxidative stress inflicted by the host immune system. To test this in our isolates, we performed *in vitro* hydrogen peroxide (H_2_O_2_) challenge assays, with the rationale that higher staphyloxanthin production would increase survival of *S. aureus* when exposed to 3% H_2_O_2_. Survival of USA300 LAC was significantly higher than the *ΔcrtN* mutant (CFU survival = 113% vs 35%; p=0.024); similarly, survival of the clinical isolates reflected their staphyloxanthin production (CFU survival = 104% SA925 vs 20% SA1088; p=0.0055) (**Figure 2F**). Growth curves (8-hour) in planktonic culture were similar across all four isolates, suggesting that differential growth rates were not responsible for this result (**Figure S2**). The survival differences between high and low staphyloxanthin producers in both clinical and USA300 LAC control strain suggest that staphyloxanthin production contributes to *in vitro* oxidative stress resistance.

To determine whether enhanced survival in SA925 and USA300 LAC was due to staphyloxanthin production as opposed to other stress responses, the isolates were grown in the presence of 100μM thymol and then exposed to 3% H_2_O_2_. This resulted in a significant drop in survival for USA300 LAC (27%; p=0.0143) and SA925 (35%; p=0.0053) (**Figure 2F**). Furthermore, when grown with thymol, their survival was no longer significantly greater than USA300 LAC *ΔcrtN* and SA1088, respectively (**Figure 2F**). These results suggest that the survival differences observed under oxidative stress are due to differences in staphyloxanthin production.

Staphyloxanthin can increase bacterial cell membrane stability by ordering the alkyl chains of membrane lipids, thus enhancing *S. aureus* survival of membrane-disrupting antibiotics (Mishra et al., 2011). To test membrane stability, we exposed *S. aureus* isolates to polymyxin B (1mM), a membrane-disrupting antibiotic. We observed a moderate decrease in survival with polymyxin B exposure in USA300 LAC *ΔcrtN* and SA1088, as compared to the staphyloxanthin producing isolates USA300 and SA925, though these findings were not statistically significant (**Figure 2G**). We thus concluded that staphyloxanthin enhances *S. aureus* survival primarily via oxidative stress resistance rather than protection from membrane disruption.

### Staphyloxanthin impairs in vivo wound healing in a diabetic murine model

To determine whether staphyloxanthin production by *S. aureus* has an *in vivo* effect on diabetic wound healing, we employed a murine model of diabetes in which mice lack the leptin receptor, *Lepr^db/db^* (referred to as db/db subsequently) (Coleman, 1978). In a pilot experiment in db/db mice, infection with high staphyloxanthin producing clinical isolate SA925 significantly delayed healing compared to low staphyloxanthin producing SA1088 or PBS control (**Figure S3A**). Guided by these preliminary studies, we designed two independent wound healing experiments in male and female 12-week-old db/db mice. Excisional 8 mm full-thickness dorsal wounds were inoculated with 2×10^8^ CFU of USA300 LAC, USA300 LAC *ΔcrtN*, SA925, SA1088 or vehicle control (PBS+10% glycerol) and covered with Tegaderm. Wounds were photographed and measured at days 0, 3, 7, and 14 in both experiments.

We compared wound closure among mice treated with *S. aureus* isolates and controls to determine whether staphyloxanthin influences diabetic wound healing. A representative image for each group and timepoint is shown in **Figure 3A**. All *S. aureus* strains impacted wound closure dynamics to some degree, but staphyloxanthin-producing strains exacerbated the delay (**Figure 3B**). At experimental day 14, wounds exposed to USA300 LAC were significantly larger than wounds exposed to USA300 LAC *ΔcrtN* (p=0.0114; **Figure 3C**). Furthermore, wounds exposed to the high staphyloxanthin-producing clinical isolate SA925 were significantly larger than wounds exposed to SA1088 (p=0.00074; **Figure 3C**). We also observed that female mice were significantly delayed in healing at 14 days post-wounding compared to male mice (p= 0.031), suggesting a sex bias in *S. aureus* infection and/or diabetic wound healing. We observed that strain SA925 caused mortality in 4 mice over the course of the 3 experiments described above, whereas no mice exposed to SA1088 died prior to the experimental endpoint (**Figure S3B**). At day 21 experimental endpoints, cultured total bacterial CFU and *S. aureus* CFU from wound tissue did not significantly differ between SA925 and SA1088 (**Figure S3C, S3D,** respectively). This suggests that delay in wound closure was not due to differences in bacterial load. Together these results indicate that increased staphyloxanthin production by *S. aureus* contributes to delayed wound healing in diabetic mice.

**Figure 3:**
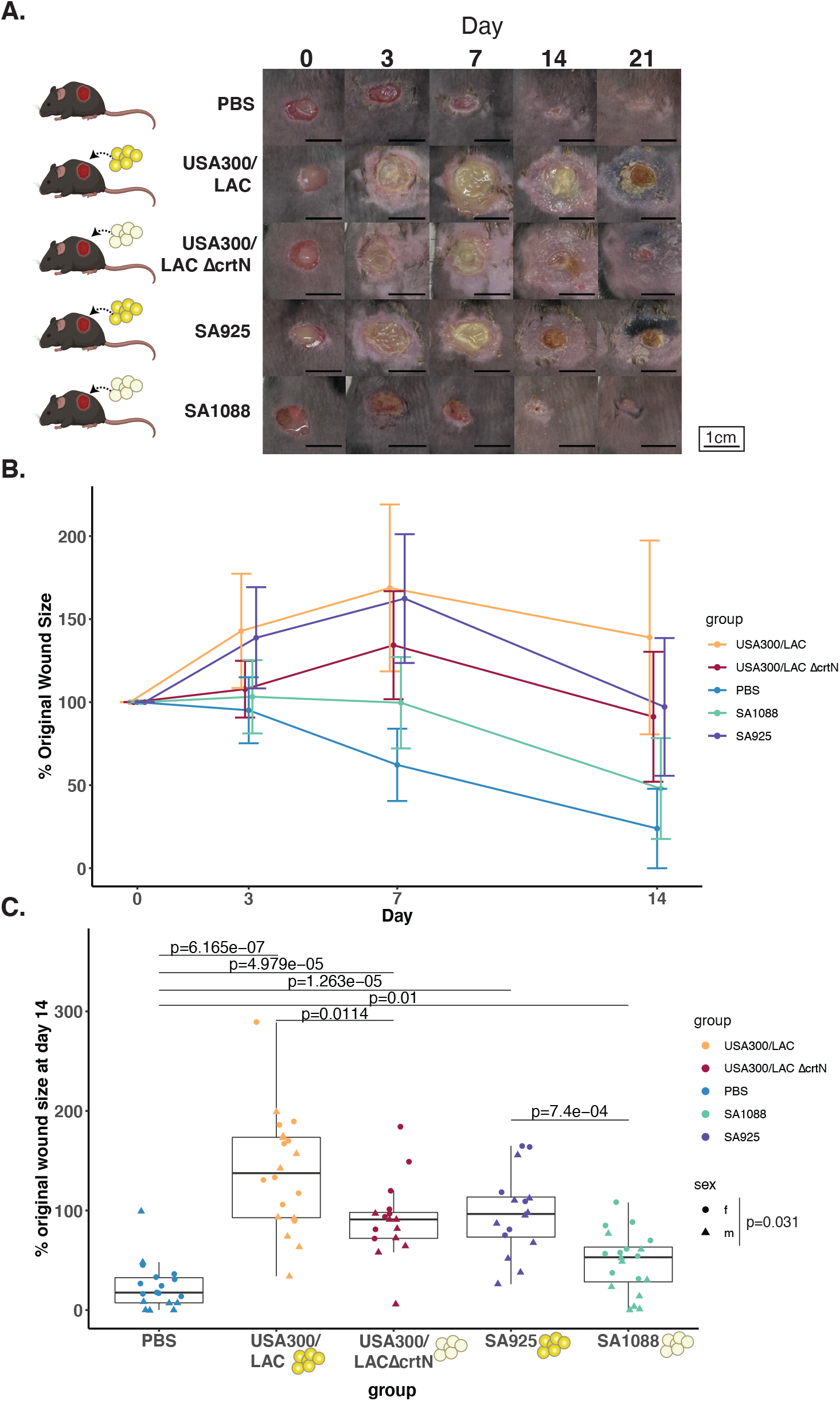
Staphyloxanthin delays wound healing in a diabetic mouse. **(A)** Full-thickness excisional wounds of male diabetic mice (db/db) were infected with 2×10^8^ *S. aureus* each, with strains USA300 LAC, USA300 *LACΔcrtN*, SA925, SA1088, or PBS control. Wound size was monitored for 21 days. Image rows follow one representative wound from each group. Scale bars are 1 cm. **(B)** Relative wound size (% original area) for day 0, 3, 7, 14 by treatment group, where dots represent the mean within the group, and error bars represent mean +/- standard deviation. **(C)** Percent change in wound size (y-axis) at day 14 post-wounding and infection by the *S. aureus* strains indicated above. Male (triangles) and female (circles) mice are indicated, where each triangle/circle represents the mean of 3 area measurements for each wound. Data represents two single sex independent experiments. A two-sided Wilcoxon rank sum test was performed amongst each group. See also Figure S3.

### Staphyloxanthin contributes to immune cell recruitment and promotes intracellular survival in neutrophils

Because staphyloxanthin is a cell surface molecule, we first hypothesized that it may contribute to neutrophil recruitment and persistence in tissue. To test this, we employed an ear wound model, where a 2 mm hole punch is created in the center of the outer ear pinna (Leung et al., 2015) in C57BL6/J mice. Punch wounds were exposed to *S. aureus* clinical isolates SA925, SA1088, or vehicle control for one week. The ear pinnae were collected and flow cytometry was performed to assess immune cell populations in the tissue (**Figure S4, Table S3**). As measured by Ly6G+ CD11b+ cells, a significantly higher percentage of neutrophils infiltrated tissue exposed to SA925 compared to both SA1088 and control (p=0.0179 and p=0.0476, respectively) (**Figure 4A; Table S3**). To determine whether these results would be consistent in the context of diabetes, we repeated the experiment with diabetic (db/db) mice. Overall, the directional trends were consistent with our observations in non-diabetic mice, though neutrophil differences between SA1088 and SA925 in db/db mice were not statistically significant (p=0.1266; **Figure 4B; Table S3**). We noted that baseline neutrophil recruitment was lower in db/db mice (6.86%) than C57BL/6 (21%), potentially due to impaired neutrophil trafficking and/or activity previously observed in db/db mice (Roy et al., 2022; Yano et al., 2012). These results together support a role for staphyloxanthin in the recruitment of neutrophils to wound tissue.

**Figure 4:**
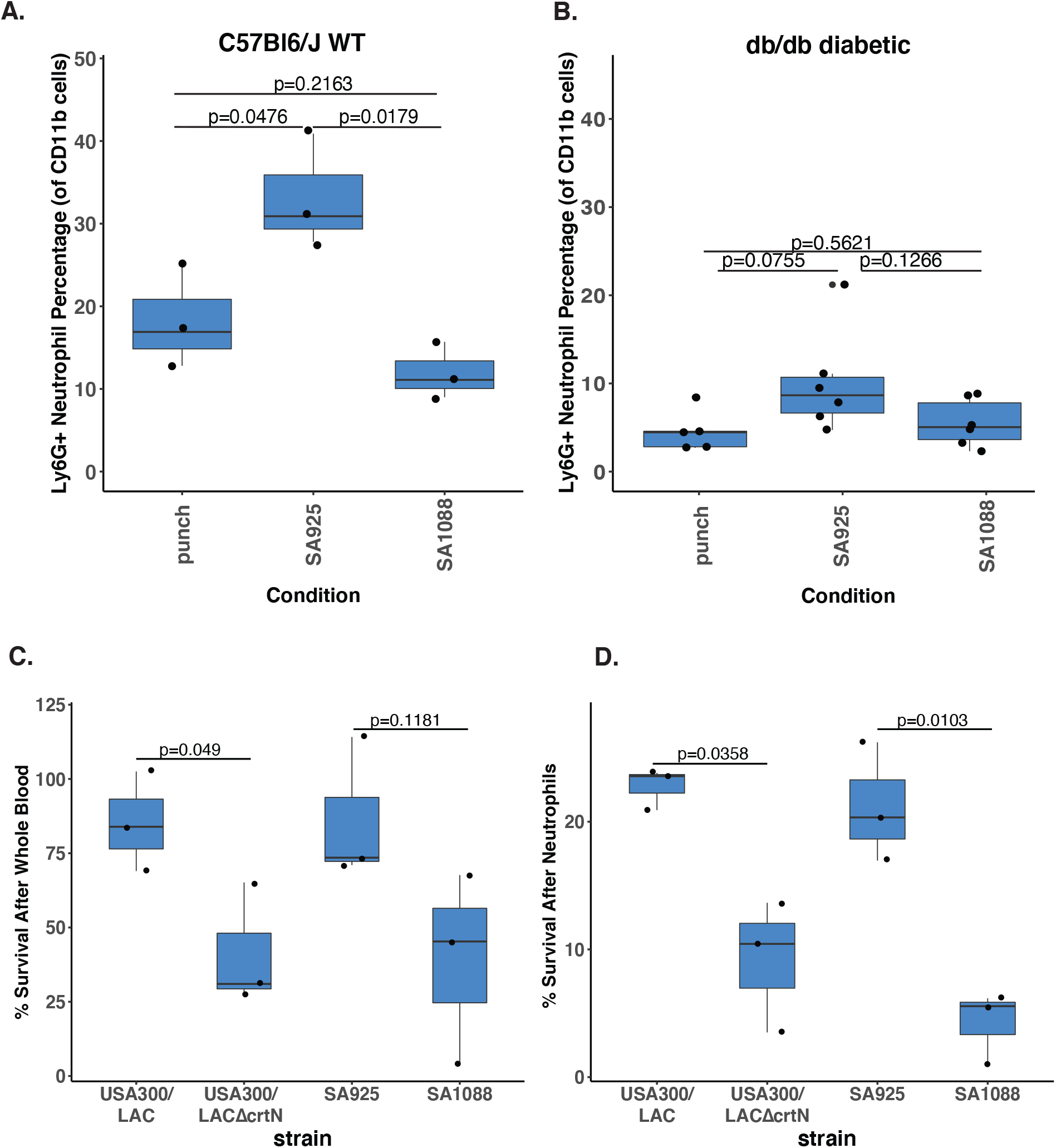
Staphyloxanthin production influences neutrophil recruitment and *S. aureus* neutrophil survival. Flow cytometry results for ear punch wounds indicating the percentage of CD11b+ cells that stained Ly6G+ from C57BL6/J **(A)** and diabetic db/db **(B)** mice infected with 2×10^8^ *S. aureus* strains. Each dot represents a mouse. Groups were compared using independent T-test. Error bars represent the standard deviation from the mean. **(C)** Survival of each *S. aureus* strain when exposed to freshly drawn whole human blood for 4 hours. **(D)** Percent survival of each *S. aureus* strain when exposed to primary human neutrophils for 30 minutes. **(C-D)** Each dot represents an independent experiment from a different donor. Independent t-tests were performed comparing each isolate. See also Table S3 and Figure S4.

Neutrophils are first line defenders against *S. aureus*, engulfing the bacteria and exposing them to reactive oxygen species via the activity of myeloperoxidase and NADPH oxidase (Beavers and Skaar, 2016). Staphyloxanthin allows *S. aureus* to survive neutrophil engulfment, thus promoting host immune evasion (Liu et al., 2005). We therefore hypothesized that the differences we observed in murine diabetic wound healing were a result of staphyloxanthin-mediated protection from neutrophil recognition and/or clearance. We first quantified survival of *S. aureus* clinical isolates and lab strains in whole human blood, an assay that measures opsonophagocytic killing by peripheral blood neutrophils. We exposed 1×10^4^ CFU of each: USA300 LAC, USA300 LAC *ΔcrtN*, SA925 and SA1088 to freshly drawn whole human blood for 4 hours before plating and quantifying CFUs. Survival of USA300 LAC was significantly greater than the staphyloxanthin mutant USA300 LAC *ΔcrtN* (p= 0.049; **Figure 4C**). Similarly, survival of SA925 was greater than SA1088 although not significantly so (p= 0.1181; **Figure 4C**). These results suggest that staphyloxanthin partially protects *S. aureus* from opsonophagocytic killing in human blood.

Therefore, we next analyzed the effects of staphyloxanthin on *S. aureus* engulfment by neutrophils. To examine the role of staphyloxanthin in *S. aureus* intracellular neutrophil survival, we performed a neutrophil survival assay by exposing freshly isolated primary human neutrophils to *S. aureus*. The neutrophils were then washed to remove any extracellular bacteria, lysed, and bacterial CFUs quantified. We observed increased survival of high staphyloxanthin producing strains USA300 LAC and SA925 compared to low staphyloxanthin producers USA300 LAC *ΔcrtN* and SA1088, (p= 0.0358 and p=0.0103 respectively) (**Figure 4D**). These results suggest that staphyloxanthin enhances intracellular *S. aureus* survival in human neutrophils.

### Whole genome comparisons between high and low staphyloxanthin isolates identify a single nucleotide variant in the *rsbU* regulator of Sigma B

To determine the genomic differences underlying the disparate staphyloxanthin phenotypes observed in the 14 isolates cultured from DFU141, we employed whole genome sequencing, comparative genomic, and phylogenetic approaches. To compare genomes at the highest structural and sequence-level resolution, we sequenced all 14 isolates using Illumina short read sequencing followed by Oxford Nanopore long read sequencing on 13/14 isolates to use as structural scaffolds for the short reads during hybrid genome assembly.

We classified the 14 assembled genomes as belonging to the CC1 lineage of *S. aureus* based on PubMLST (Jolley et al., 2018). We annotated each genome using Prokka (Seemann, 2014), and estimated their core and accessory genomes using Roary (Page et al., 2015) including the CC1 MW2 reference genome as a closely related outgroup. An estimated maximum likelihood phylogeny grouped the three low staphyloxanthin producing isolates together in a monophyletic clade (**Figure 5A**), suggesting this genotype/phenotype combination may have arisen intra-host. We performed pairwise alignments and comparisons of the 14 genomes for structural variants, single nucleotide polymorphisms (SNPs), and large scale insertions/deletions using NucDiff (Khelik et al., 2017). These comparisons identified exactly three SNPs and one 9-base deletion which were consistently present in low staphyloxanthin genomes but absent from high staphyloxanthin genomes. Of these four consistent variants, one SNP and the deletion (**Figure 5A, Panels ii and iii**) were predicted by both Prokka and PGAP (Haft et al., 2018) annotations to be in noncoding regions. One SNP (**Figure 5A, Panel i**) corresponding to a Thr→Ala amino acid change was predicted by Prokka to be in a noncoding region between *folK* and *LysS*, but predicted by PGAP to be in a hypothetical ORF between the two genes. Another SNP (**Figure 5A, Panel iv**) was located in the positive Sigma B regulator *rsbU*. This corresponded to a Met→Leu amino acid change in the active phosphatase domain of *rsbU*.

**Figure 5:**
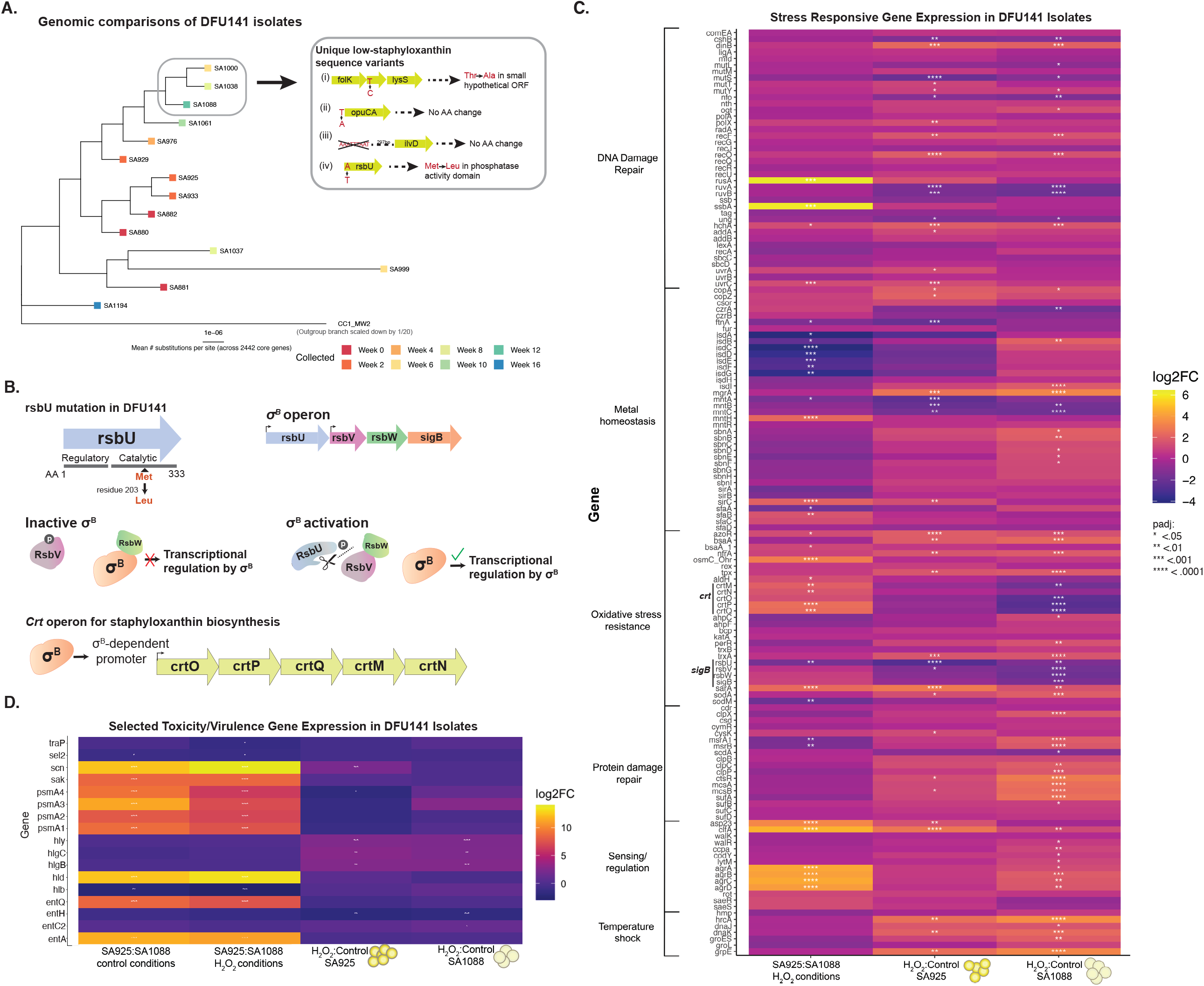
Genetic and transcriptomic analysis of high and low staphyloxanthin-producing clinical isolates. **(A)** Maximum likelihood phylogeny of all isolates collected from DFU141. Tree nodes are colored by the week collected, and staphyloxanthin-deficient isolates are circled. The inset box depicts the four unique sequence variants shared by all three and their predicted amino acid changes. **(B)** A single nucleotide change was identified in *rsbU* as the likely cause of low staphyloxanthin production in SA1088. RsbU is a regulator and part of the Sigma B operon. Sigma B regulates staphyloxanthin biosynthesis. **(C)** Expression of selected virulence and toxicity genes (y-axis) in SA925 and SA1088 with and without 1% hydrogen peroxide (H_2_O_2_; x-axis). **(D)** Expression of selected stress response genes (y-axis) of SA1088 and SA925 with and without H_2_O_2_ (x-axis). **(C-D)** * depicts log2Fold change p<0.05. See also Figure S5.

The *rsbU* gene encodes a well-characterized phosphatase regulator of the Sigma B operon which, in response to environmental stress, dephosphorylates RsbV which prevents inhibition of Sigma B by RsbW (**Figure 5B**) (Palma and Cheung, 2001). The transcription factor Sigma B, in turn, positively regulates the *crt* biosynthetic operon for staphyloxanthin. Along with the fact that this non-synonymous SNP corresponds to an amino acid change in the catalytic domain of the protein, the *rsbU* variant is most likely responsible for impaired staphyloxanthin production in DFU141 isolates. Consistent with previous studies (Giachino et al., 2001), we found that *rsbU* was necessary for staphyloxanthin production (**Figure S5**). Still, this variant represents a change in protein sequence which may alter but not necessarily abolish RsbU’s regulatory function. This motivated our subsequent investigation into the transcriptomic differences between high- and low-staphyloxanthin DFU141 isolates, and how the two strains respond to oxidative stress at the transcriptional level.

Though the four genetic variants summarized in **Figure 5A** were the only ones consistently present in the 3 low staphyloxanthin-producing isolates and absent from the 11 high staphyloxanthin-producing isolates, there were other pairwise differences among isolates. For example, the representative staphyloxanthin-producing isolate SA925 contains a phage insertion absent from SA1088 with >99% nucleotide identity to the □Sa3mw phage from *S. aureus* MW2 (Genbank BA000033.2) which contains *sak, scn*, and staphylococcal enterotoxins a and q (*entQ* and *entA*), among other phage-associated genes (Baba et al., 2002).

### Staphyloxanthin alters expression of stress response genes under oxidative stress conditions and protects *S. aureus* from oxidative damage

We investigated transcriptional differences between the high and low staphyloxanthin producing isolates from DFU141 when exposed to oxidative conditions. To evaluate this, we conducted RNAseq analysis on SA925 and SA1088 with and without 1% H_2_O_2_ exposure. Under H_2_O_2_ exposure, expression of genes in the *crt* operon was significantly higher in SA925 compared to SA1088 (**Figure 5C**). Consistent with previous reports, (Palazzolo-Ballance et al., 2008; Ramond et al., 2021), under H_2_O_2_ stress, *sigB* operon genes were generally down regulated in both strains, but more significantly so in SA1088. The exception was *rsbU*, which was significantly more downregulated in SA925 compared to SA1088 (**Figure 5C**). Quorum sensing operon *agrABCD* was more highly expressed in SA925 compared to SA1088 under H_2_O_2_ conditions. Interestingly, *agrABCD* was upregulated in SA1088 in response to H_2_O_2_. This coincided with H_2_O_2_-responsive downregulation of *rsbU* in SA1088, suggesting that while SA1088’s mutant *rsbU* may be incapable of adequate *sigB* activation to promote *crt* expression, it may still be able to downregulate *agr* as previously reported (Olivier et al., 2009).

We hypothesized that stress response genes, and in particular those responsible for mitigating oxidative stress, would be upregulated in response to H_2_O_2_. Indeed, we observed that the expression of a number of stress response genes increased significantly under these conditions in both isolates including: *sodA, tpx, trxA*, and *nfrA, sarA* (**Figure 5C**). Intriguingly, we found that several oxidative stress sensing and resistance genes were significantly upregulated in SA1088, but not SA925, including *ahpC, perR, lytM, and walR*. These genes could offer compensatory resistance mechanisms to the oxidative stress responses impeded by the *rsbU/sigB* defect in SA1088. Both strains exhibited changes in expression of DNA damage repair genes in response to H_2_O_2_, though SA1088 uniquely upregulated the endonuclease *nth* while SA925 uniquely upregulated *polX, addA*, and *uvrA*. Strikingly, SA1088 upregulated more genes related to damaged protein repair and degradation, including the *clpC* operon, *msrAB*, and *sufAB* (Stahlhut et al., 2017; Wozniak et al., 2012). This suggests that SA1088, deficient in staphyloxanthin and other stress resistance traits, may incur more damage to its proteins by H_2_O_2_, necessitating the activation of these protein repair and degradation pathways.

Iron-related genes were also more upregulated in SA1088 compared to SA925 under oxidative conditions. The iron surface determinant (*isd*) genes were almost all more highly expressed in SA1088 than SA925 in the presence of H_2_O_2_. Furthermore, *isdB, isdI, mgrA* and multiple staphyloferrin B (*sbn*) operon genes were uniquely upregulated in SA1088 in response to H_2_O_2_ exposure (**Figure 5C**).

In addition to differing stress responses, we observed several toxicity-associated genes to be expressed more highly in SA925 compared to SA1088 under both conditions, some of which likely result from the presence of the MW2 phage in SA925 (**Figure 5D**). Interestingly, though enterotoxin A (*entA*), enterotoxin Q (entQ), staphylokinase (*sak*), and staphylococcal complement inhibitor n (*scn*), are all found on the MW2 phage and therefore missing from SA1088, only *scn* was upregulated in SA925 in response to H_2_O_2_ exposure. Other toxicity genes not found on the MW2 phage insertion were also differentially expressed between the strains. Phenol-soluble modulin components (*psmα1-4*) and delta hemolysin (*hld*), for example, were upregulated in SA925 compared to SA1088 under both conditions despite the genes being present in both strains, but none of these genes showed altered expression in response to H_2_O_2_ (**Figure 5D**).

Overall, there were more stress response genes upregulated in SA1088 compared to SA925 under H_2_O_2_ conditions, particularly those related to iron uptake and storage, damaged protein repair and degradation, and oxidative stress mechanisms other than staphyloxanthin. These findings suggest that the staphyloxanthin-deficient SA1088 experiences greater stress in response to H_2_O_2_ than the staphyloxanthin-competent SA925. This is consistent with our findings that high staphyloxanthin strains experience greater H_2_O_2_ survival and suggests that staphyloxanthin embedded in the *S. aureus* cellular membrane protects it from oxidative stress.

## DISCUSSION

*S. aureus* is a common pathogen to injured skin and wounds, and particular strains of *S. aureus* have been shown to delay diabetic wound healing. Nevertheless, because the effects of *S. aureus* are strain-specific, it is unclear which *S. aureus* phenotypes contribute to poor healing and infection-related wound outcomes. In this work, we sought to identify *S. aureus* genetic factors associated with poor healing outcomes. To accomplish this goal, we performed high-throughput phenotyping screens on a collection of 221 *S. aureus* isolates from 60 DFU. We discovered that production of the pigment staphyloxanthin, but not siderophore, staphylokinase, or biofilm production, was significantly associated with delayed healing (**Figure 1**).

While our results suggest that *S. aureus in vitro* biofilm formation, staphylokinase and siderophore production are not associated with a delayed healing outcome, it does not necessarily mean that these phenotypes do not play a role in the context of diabetic wounds. There are several possible explanations. For example, *in vitro* biofilm formation of *S. aureus* alone may not delay wound healing, but wounds are often communities consisting of multiple species and strains of bacteria as well as fungi (Kalan et al., 2019). Some of these bacteria such as *Pseudomonas aeruginosa* are well known to form robust biofilms (Lee and Yoon, 2017). Therefore, one possibility is that *S. aureus* does not need to form a strong biofilm because other species are producing biofilms which inhibit healing and facilitate a hospitable environment for *S. aureus*. We unfortunately also cannot rule out the possibility that while *S. aureus in vitro* biofilm formation is not associated with delayed healing, it is possible that there are additional host, microbiome, or other environmental factors that change the biofilm dynamics *in vivo*. Further study into *S. aureus in vivo* biofilms will be required before they can be adequately modeled by a high-throughput screen such as ours.

Beyond studying staphyloxanthin in a high throughput assay, we also wanted to understand the role of staphyloxanthin in DFU clinical isolates. Having found that DFU141 exhibited dichotomous staphyloxanthin phenotypes with either high and low staphyloxanthin-producing isolates, we selected two of these isolates, SA925 (high) and SA1088 (low) to study further. This gave us the opportunity to closely examine host-microbe interactions in two genetically similar clinical isolates with distinct staphyloxanthin phenotypes. However, these clinical isolates contained several differences including a mutation in an upstream stress operon (**Figure 5**). Parallel comparisons between USA300 LAC and its mutant USA300 LAC *ΔcrtN* allowed us to isolate the effects of staphyloxanthin production. From our work with these four isolates *in vitro*, we were able to verify previous findings that staphyloxanthin enhances hydrogen peroxide survival, modestly improves polymyxin survival, and that thymol abrogates the staphyloxanthin-mediated H_2_O_2_ survival (**Figure 2**).

Consistent with our observed association between staphyloxanthin production and delayed healing outcome *in vitro*, we demonstrated that in a diabetic mouse model, wounds exposed to high-staphyloxanthin *S. aureus* isolates were significantly larger and more severe than those exposed to low staphyloxanthin *S. aureus* isolates (**Figure 3**). Furthermore, our work supports a neutrophil-based mechanism for this effect. High-staphyloxanthin producing isolates survived better when engulfed by neutrophils or exposed to whole human blood (**Figure 4C,D**). This suggests that the ability of staphyloxanthin to shield *S. aureus* from reactive oxygen species produced by the host immune system is enabling it to persist and thrive in the wound. This could explain why high-staphyloxanthin exposed wounds display a delayed healing phenotype. Interestingly, we also observed that SA925 recruited significantly more neutrophils in C57BL/6 compared to SA1088. We observed the same directional trend in db/db mice, though it was not significantly different (**Figures 4A,B**). This may be because neutrophil recruitment is lower overall in diabetic patients (Roy et al., 2022). Nonetheless, these findings may suggest that higher staphyloxanthin production stimulates increased neutrophil recruitment. This would promote a prolonged inflammatory phenotype that is consistent with delayed wound healing. In our RNAseq analysis we observed enhanced expression of cell wall components including capsule polysaccharides in SA925 as compared to SA1088 (**Figure 5C**). We suspect that enhanced expression of neutrophil attracting cell wall components may explain why higher staphyloxanthin strains appear to attract more neutrophils in our experiments.

Despite the genetic similarity between SA925 and SA1088, RNAseq analysis revealed substantial gene expression differences, with and without exposure to H_2_O_2_. SA1088 upregulated more genes involved in iron transport and storage under H_2_O_2_ stress compared to SA925 (**Figure 5C**). This is consistent with a previous study demonstrating that a *rsbU* mutant form of the MW2 strain upregulated iron storage and transport genes in response to H_2_O_2_ stress (Palazzolo-Ballance et al., 2008). Due to its tendency to undergo Fenton chemistry, iron is a known catalyst of oxidative stress, and it has been previously suggested that *S. aureus* sequesters iron to protect itself (Touati, 2000). We hypothesize that iron transport and storage genes are only upregulated in SA1088 because the staphyloxanthin in SA925 neutralizes some of the excess free radicals protecting the bacteria from the deleterious effects of iron in the presence of excess H_2_O_2_. Simultaneously, SA1088 upregulated many genes responsible for repairing stress-induced protein damage and targeting damaged proteins for degradation (**Figure 5C**), suggesting that SA1088 is more vulnerable to oxidative damage by H_2_O_2_ than SA925. Collectively, these findings support our hypothesis that high staphyloxanthin production protects *S. aureus* from oxidative stress.

Under H_2_O_2_ stress, *rsbU* expression was lower in SA925 compared to SA1088, in which *rsbU* is mutated. We hypothesize that SA1088 expresses more *rsbU* than SA925 to compensate for lost functionality due to its mutation in the phosphatase domain. This hypothesis is supported by our finding that the *agr* quorum sensing system is significantly upregulated in SA925 compared to SA1088, and that downregulation of the *sigB* operon in SA1088 in response to H_2_O_2_ coincides with increased *agr* expression. Previously, it has been shown that RsbU inhibits *agr* expression (Bischoff et al., 2001). Our findings that *agr* is expressed more highly in SA925 suggests that excess compensatory RsbU is strongly downregulating *agr* expression in SA1088. Perhaps because the expression of the *agr* system was upregulated in SA925, phenol soluble modulins (psms) were dramatically upregulated in SA925 compared to SA1088. Previously, psms have been shown to be responsible for lysing neutrophils after phagocytosis, permitting *S. aureus* to escape (Surewaard et al., 2013). This difference between SA925 and SA1088 may contribute to the enhanced survival of SA925 in human neutrophils. However, because we observed significantly reduced survival of USA300 LAC *ΔcrtN* in neutrophils and under H_2_O_2_ stress compared to USA300 LAC, staphyloxanthin must play a direct role as well (**Figure 4**).

Overall we found that staphyloxanthin promotes resistance to oxidative stress. However, it is possible that staphyloxanthin contributes to delayed healing in a manner beyond oxidative stress resistance. Previously, it has been demonstrated that staphyloxanthin reduces cell-membrane fluidity, thus partially protecting *S. aureus* from some chemical assaults (Kossakowska-Zwierucho et al., 2016; Mishra et al., 2011). We observed that staphyloxanthin producing isolates only modestly enhanced polymyxin B survival (**Figure 2G**), in line with previous studies (Mishra et al., 2011). The greatest effects of staphyloxanthin that we observed were on neutrophil survival and oxidative stress resistance (**Figures 2F,4D**). These findings suggest that staphyloxanthin promotes resistance to oxidative stress, which plays a substantial role in the interaction between neutrophils and *S. aureus*.

In our current study we identified staphyloxanthin as a *S. aureus* phenotype associated with delayed healing in DFU. There were several limitations, including the recalcitrant nature of *S. aureus* clinical isolates for genetic manipulation, limiting our ability to create staphyloxanthin knockouts in the parent strains. We were also unable to directly measure staphyloxanthin production in the wound, though such approaches could have clinical utility for directly assessing a potential risk factor for complication. Novel strategies to manage and treat DFU, especially those with low risk of further promoting anti-microbial resistance, are urgently needed. Our study puts forth staphyloxanthin as a promising new target for DFU colonized or infected with *S. aureus*.

## ACKNOWLEDGEMENTS

We thank Ms. Jamie Pan and Ms. Anisa Ray for their assistance with phlebotomy; Dr. Paul Planet for his generous gift of *S. aureus* strains 502A and JE2 (USA300 LAC); Dr. Daniel Shin, Associate Director of the Data Science and Informatics Core of the Penn SBDRC for statistical consultation. The following reagent was provided by the Network on Antimicrobial Resistance in Staphylococcus aureus (NARSA) for distribution by BEI Resources, NIAID, NIH: Staphylococcus aureus subsp. aureus, Strain JE2, Transposon Mutant NE382 (SAUSA300_2498), NR-46925. We thank current and former members of the Grice lab and the Department of Dermatology for critical discussion and review of the work. This work was funded by the following grants to EAG from the NIH, NIAMS (R01AR006663), NINR (R01NR015639 to EAG and R01NR009448 to SEG), the NIDDK Diabetic Complications Consortium (DiaComp, www.diacomp.org, grant DK076169), the Burroughs Wellcome Fund PATH Award, and the Dermatology Foundation Sun Pharma Research Award. This research was also supported by the Penn Skin Biology and Disease Resource-based Center (Penn SBDRC supported by NIH/NIAMS P30AR069589). ARM-V was supported by the Penn Dermatology Research T32 Training Grant (NIH/NIAMS T32AR007465); AEC was supported by the UPenn Blavatnik Family Fellowship; AU is supported by the Prevent Cancer Foundation Awesome Games Done Quick fellowship; EKW is supported by NIAMS F31AR079852; VML is supported by NIAMS F31AR079901. Figure 1A was created with Biorender.com

## METHODS

### EXPERIMENTAL MODEL AND SUBJECT DETAILS

#### Animal models and husbandry conditions

All mouse experiments were conducted under protocols approved by the University of Pennsylvania Institutional Animal Care and Use Committee (Protocol 804065). All mice were housed and maintained in an ABSL II and specific pathogen free facility in the Clinical Research Building vivarium at the University of Pennsylvania. The following strains of mice were used in these studies: C57BL6/J (JAX stock #000664) and B6.BKS(D)-*Lepr^db^*/J (JAX stock #000697)(Coleman, 1978).

##### Excisional wound model

Both male (n=24) and female (n=33) mice were used and housed individually during 3 independent experiments. Dorsal region was shaved when mice were 12 weeks old. Two full thickness excisional wounds of 8mm were created by punch Biopsy tool (Miltek) on shaved dorsal skin. Wounds were inoculated wtih 2 x10^8^ CFU of *S. aureus* or PBS with 10% glycerol vehicle control and the wound covered with transparent film (Tegaderm, 3M). Each wound was photographed and measured at the time of wounding (t=0), day 3, day 7 and day 14. For experiment #3 with males, wounds were also photographed and measured at day 21. After day 7, the transparent film was removed. All experimental procedures were conducted on mice anesthetized with isoflurane and Bupivacaine (0.1 mg) and Bupinorphine-SR (0.05mg) were administered for pain relief. Mice were administered Diet Gel Recovery (Clear H_2_O) every 3 days to prevent weight loss. Wound measurements were made in ImageJ and represent the mean of 3 independent measurements. Wilcoxon sum rank tests were performed to compare groups.

##### Ear punch wound model

C57BL/6 or db/db mice were ear-punched using a 2 mm Animal Ear-Punch (Fisher Scientific) and 1×10^8^ CFU of bacteria topically applied to the entire mouse using sterile cotton swabs at Day 0 and Day 4 post ear-punch. At Day 7 post ear-punch, pictures of the punch were taken for calculation of wound area and a digital caliper (Model PK-0505, Mitutoyo) was used to measure ear thickness. Additionally, ear tissue was collected for CFU calculation and flow cytometry.

#### Human blood and neutrophils

Primary human neutrophils and whole blood were freshly drawn from healthy human subjects according to a protocol approved by the University of Pennsylvania Institutional Review Board #851659. Both male and female subjects between ages 28 and 61 were used. 3 individuals were used in each experiment.

#### Microbial Strains

##### DFU100 *S. aureus* strains

As previously described in Gardner et al. 2014, swab specimens were collected from 100 DFU during the course of a prospective cohort study of wound bioburden and clinical outcomes. Swabs were collected using the Levine technique to sample deep tissue fluid (Gardner et al., 2006). The swabs were placed in charcoal transport media and transported to the microbiology laboratory, where they were then vortexed in 1 mL of tryptic soy broth prior to dilution and plating. *S. aureus* was identified on Columbia blood agar (Remel) as yellow beta-hemolytic colonies that stained as Gram-positive cocci and tested catalase positive as well as *Staphylococcus* latex-agglutination positive. Unique isolates were banked as glycerol stocks and stored at −80C.

##### Reference S. aureus strains

We used the following laboratory reference strains: *S. aureus* strains SA113 (ATCC #35556), 502A (gift of Paul Planet), JE2 (gift of Paul Planet). The following reagents were provided by the Network on Antimicrobial Resistance in Staphylococcus aureus (NARSA) for distribution by BEI Resources, NIAID, NIH: *Staphylococcus* aureus subsp. aureus, Strain JE2 (Paudel et al., 2021), NR-46543, and the corresponding transposon mutants:

NE382 (SAUSA300_2498), NR-46925
NE1607 (SAUSA300_2025), NR-48149
NE1109 (SAUSA300_2022), NR-47652
NE1472 (SAUSA300_2023), NR-48014
NE1872(SAUSA300_2024), NR-48414

### METHOD DETAILS

#### *In vitro* Biofilm Production

Clear 96-well plates were coated with bovine fibronectin (Millipore Sigma) by diluting 1 mg/mL bovine fibronectin solution into PBS to a final concentration of 25 μg/mL and adding 60 μL per well, then incubating coated plates for 1 hour at 37° C. Excess fibronectin was removed by aspiration and plates washed 2 times with PBS. Overnight cultures of *S. aureus* were grown in trypticase soy broth (TSB) from single colonies and then diluted 1/100 in TSB with 5% dextrose and arrayed in technical triplicate. Each plate was then grown statically at 37° C for 24 hours to allow biofilm matrix to form. Bacterial growth was visualized by plate reader at OD600 nm. The media was removed and the plate was gently washed by dipping it in distilled water. The biofilm was stained with 125 μL of 0.5% crystal violet solution (Millipore Sigma) in water and the plate was incubated for 5 minutes. 200uL of acetic acid was added to each well to solubilize crystal violet and pipetted to mix, then quantified on a Biotek Synergy HT plate reader at OD540 nm. Three biological replicates were performed for each isolate. For normalization, each biofilm was compared to production from *S. aureus* SA113 grown on the same plate. Percent biofilm production was calculated by dividing mean OD540/OD600 readings for each isolate and comparing to SA113 biofilm production.

#### Staphyloxanthin production

Cultures of each *S. aureus* isolate were grown in 5 mL TSB shaking at 37° C for 24 hours from single colonies. Overnight growth was quantified on a Biotek Synergy HT plate reader at 600nm. Cultures were centrifuged for 1 minute at 13,000 RPM and supernatant was removed. Each pellet was resuspended in 700 μL of methanol and incubated at 55° C for 10 minutes. Tubes were centrifuged for 2 minutes at 13,000 RPM. 300 μL from each tube was added to each well of a clear 96-well plate in triplicate and then quantified at OD465 nm on a Biotek Synergy HT plate reader. Three biological replicates were performed for each isolate. For normalization, all isolates were compared to staphyloxanthin production from *S. aureus* 502A strain on the same plate. Percent staphyloxanthin production was calculated by dividing mean OD540/OD600 readings for each isolate and comparing to 502A staphyloxanthin production.

#### Staphylokinase production

Overnight cultures of each *S. aureus* isolate were grown in TSB shaking at 37 °C from single colonies. An aliquot of each culture (100 μL) was diluted 1:1 in PBS and OD600 was quantified on a Biotek Synergy HT plate reader. Simultaneously 1 mL was removed from each culture and centrifuged at 13,000 RPM for 1 minute. Three 100μL aliquots of supernatant were removed to 3 wells on a clear 96-well plate. To the supernatants, 50 μL of human glu-plasminogen (0.04mg/mL; Invitrogen) in Tris buffered saline was added to each well and mixed by pipetting. The plate was incubated for 15 minutes at 37 °C, followed by the addition of 20 μL plasmin specific chromogenic substrate S-2251(H-D-Val-Leu-Lys-paranitroanilide, 2 mmol/L, Chromogenix) to each well. Color was allowed to develop for 3 hours at 37 °C then measured at OD405 on a plate reader. Color was compared to a standard curve of purified staphylokinase diluted 2x from 2μg to 0.0625 μg.

#### Siderophore production

Using a protocol adapted from (Schwyn and Neilands, 1987), overnight cultures of each *S. aureus* isolate were grown from single colonies in TSB with shaking at 37 °C. An aliquot of 100 μL from each culture was diluted 1:1 with PBS and OD600 was quantified on a Biotek Synergy HT plate reader. One OD600 of cells was removed from each culture to an Eppendorf tube and centrifuged at 13,000 rpm for 1 minute. Supernatant was removed and cells were resuspended in 1mL of PBS and incubated at 37 °C for 48 hours. Shuttle solution was added to previously described CAS solution. Bacterial tubes were centrifuged for 1 minute at 13,000 rpm. An aliquot of 500 μL of supernatant from each tube was added to a pre-prepared Eppendorf tube containing 1mL of CAS solution. Tubes were incubated for 2 hours at room temperature in the dark. An aliquot of 300 μL from each tube was added to a clear 96 well plate in triplicate. Siderophore production was quantified on a plate reader at OD630. Three biological replicates were performed for each isolate.

#### Hydrogen peroxide survival assays

Cultures of *S. aureus* were grown in TSB overnight from single colonies. 500 μL of each culture was centrifuged at 13000 rpm for 1 minute. Supernatant was removed and cellular pellet was resuspended in 1 mL of PBS. 450 μL from each resuspended culture was aliquoted into two Eppendorf tubes. To one aliquot, an additional 450 μL of PBS was added. To the second aliquot, 450 μL diluted hydrogen peroxide was added. To dilute the hydrogen peroxide, 2 mL of stabilized 30% H_2_O_2_ (Thermo-Fisher) was added to 7 mL of water and mixed. The cultures were incubated for 2 hours at 37° C then plated onto blood agar plates to quantify CFUs. Survival was calculated by comparing the plates with and without H_2_O_2_ for each culture.

#### Polymyxin B survival assay

Cultures of *S. aureus* were grown in TSB overnight from single colonies. An aliquot of 200 μL of each culture was diluted 10x into 1x PBS and then split into two separate tubes. To one tube, 10 μL of 100 mM polymyxin B in ethanol was added. To the other tube 10 μL of ethanol was added. Tubes were incubated at 37° C for 30 minutes with shaking and then each tube was plated on blood agar and CFUs quantified to compare survival with and without polymyxin.

#### Antibodies and flow cytometry

Single cell suspensions from the ear were obtained by using tweezers to separate the dorsal and ventral sides of the mouse ear and placing the sheets dorsal-side down in a 24 well plate with 1 mL/well of RPMI1640 containing 0.25 mg/mL of Liberase TL (Roche, Diagnostics Corp.) and 10 μg/mL of DNase I (Sigma-Aldrich). Sheets were incubated for 90 minutes at 37° C and a single cell suspension was obtained by dissociation using the plunger of a 3 mL syringe to break up the tissue on a 40 μm cell strainer (Falcon). Dissociated tissue was washed into a 50 mL conical using PBS containing 0.05% BSA and 20 μM EDTA. Staining for surface markers was performed by incubating the cells in single cell suspension with anti-mouse CD16/CD32 mouse Fc block (Thermo Fisher Scientific, RRID: AB_467135) and 10% rat-IgG in PBS. Dead cells were stained using LIVE/DEAD Fixable Aqua Dead Cell Stain Kit (Thermo Fisher Scientific, L34957) and surface markers were stained using CD45 (Clone: 30-F11, BioLegend, Cat 103108), CD8b (clone: YTS156.7.7, BioLegend, Cat 126609), CD11b (clone: M1/70, BioLegend, cat 101235), CD90.2 (clone: 30-H12, BioLegend, cat 105343), Ly6C (clone: HK1.4, BioLegend, cat 128041), Ly6G (clone: 18A, BioLegend, cat 127614), γδ TCR (clone: GL3, BioLegend, cat 118136), F4/80 (clone: t45-2342, BD Pharmigen, cat 565410), CD4 (Thermo Fisher Scientific, cat MCD0417), and CD11c (clone: N418, BioLegend, cat 117318). Cells were fixed using 2% formaldehyde and the data was collected using the LSR Fortessa flow cytometer (BD) and analyzed using FlowJo Software (BD).

#### Whole blood survival assay

Freshly drawn whole human blood (25 μL) was mixed with 1×10^4^ *S. aureus* and incubated with shaking for 4 hours at 37° C. Samples were plated on blood agar and CFU quantified.

#### Neutrophil survival assay

Primary human neutrophils were isolated from freshly drawn human blood using Neutrophil Isolation Medium. Remnant red blood cells were removed from neutrophil pellet using RBC lysis buffer. *S. aureus* (4.5×10^6^ CFU) was pre-incubated with 10% human serum for 15 minutes, washed with PBS, added to 3×10^5^ neutrophils, and incubated at 37° C for 30 minutes. Gentamicin (400 μg/mL) was added to each tube to kill extracellular *S. aureus*. Pelleted neutrophils were washed with PBS then lysed with water (pH = 11.0). Neutrophil lysates were plated onto blood agar and bacterial CFUs were counted.

#### Whole genome sequencing

All DNA extractions were performed on overnight cultures of DFU141 isolates grown in 5 mL TSB from single colonies. Genomic DNA extractions for short read sequencing were prepared using the *Quick-DNA* Miniprep Plus Kit (Zymo Research). High molecular weight genomic DNA extractions were prepared using the Monarch HMW DNA Extraction Kit for Tissue (NEB). All 14 isolates cultured from DFU141 were initially sequenced with paired-end sequencing on Illumina HiSeq 2500 at PennCHOP Microbiome Core. SA929 and SA882 were resequenced on NextSeq 2000 at the Microbial Genome Sequencing Center (MiGS) due to suspected contamination (see Whole Genome Assembly & Quality Control). All 14 isolates except DOERN929 were further sequenced using Oxford Nanopore long read sequencing at MiGS.

#### Whole genome assembly, processing, and quality control

Raw, demultiplexed short reads were trimmed using TrimGalore [https://www.bioinformatics.babraham.ac.uk/projects/trim_galore/] based on the output of FastQC[https://github.com/s-andrews/FastQC]. Long reads were trimmed using Porechopper [https://github.com/rrwick/Porechop]. Hybrid assembly was performed on the Illumina & Oxford Nanopore-sequenced isolates using Unicycler (Wick et al., 2017) with default “Normal” settings for hybrid assembly and Illumina short read-only assembly, respectively. SA929 was assembled using Unicycler’s wrapper for SPAdes *de novo* assembly, and contigs <100 bp in length were removed. We performed Mash screens against each assembled genome to confirm its identity as *Staphylococcus aureus* (Ondov et al., 2019). To check for intra-species contamination in the short reads, trimmed short reads from each isolate were mapped to a SPAdes *de novo* assembly for that isolate and SNP-called using Bowtie2 and Samtools, respectively (Langmead and Salzberg, 2012; Li, 2011). Based on Raven et al.’s 2020 benchmarking study in clinical MRSA sequences, we counted the number of SNPs in each alignment with ≥10X depth located at least 50 bp apart from one another, and considered assemblies contaminated if they contained >30 such SNPs. DOERN882 and DOERN929 were re-sequenced and successfully re-checked for contamination based on this metric. To check for contamination in hybrid genome assemblies, we performed HMM searches for Anvio’s set of bacterial universal single copy genes (USCGs) in each assembled genome and screened genomes for multiple copies of theoretically single copy genes.

#### Phylogeny Estimation

PubMLST assigned each of the 14 *S. aureus* isolates from DFU141 to Clonal Complex 1 (CC1). We therefore estimated a maximum likelihood phylogeny from the 14 *S. aureus* genomes from DFU141 along with a CC1 MW2 strain reference genome outgroup (NCBI accession *GCF_000011265.1*). We annotated CDS loci in each genome assembly using Prokka (Seemann, 2014), and performed a probabilistic multiple alignment of shared (core) genes using Roary and PRANK (Löytynoja and Goldman, 2010; Page et al., 2015a). A maximum likelihood phylogeny was estimated from the core gene alignment using RaxML’s GTRGAMMA model with 100 random starts (Stamatakis, 2014).

#### Genomic Analysis of DFU141 Staphyloxanthin Mutants

To identify variants consistently and uniquely found in the three low staphyloxanthin-producing genomes from DFU141 as compared to the 11 high-staphyloxanthin-producing genomes, we first annotated 43 genomic differences between SA925 (high) and SA1088 (low) using NucDiff (Khelik et al., 2017). We then annotated the presence/absence of each of these 43 variants in comparisons of SA925 to each of the other 12 genomes from DFU141, identifying 4 variants total which were consistently found in comparisons of SA925 to all three low-staphyloxanthin isolates, but none of the high-staphyloxanthin isolates. We further characterized each of these 4 variants by their genomic position relative to predicted coding regions in annotations by both Prokka and PGAP (Haft et al., 2017). For variants predicted to be in coding regions, we performed BLAST alignments of predicted amino acid sequences for the CDS containing each variant from SA925 and SA1088 [Supplementary Table containing full list of variants].

#### Bacterial RNA sequencing

Cultures of SA925 and SA1088 (5 mL) were grown overnight in TSB from single colonies. Samples were back diluted 1/1000 and grown with and without 1% H_2_O_2_ shaking at 37° C to an OD 600 of 1.0. RNA was extracted from each sample by resuspending each bacterial pellet in 1 mL TriZol and homogenizing in a BioSpec MiniBeadbeater 16 with 0.1 mm zirconium particles (BioSpec) followed by centrifuging lysate at 14,000 g for 3 minutes. Following centrifugation, supernatant was mixed with .5mL 95% ethanol and treated using the Direct-zol RNA Miniprep Plus Kit (Zymo Research) and treated with DNAse 1. RNA integrity was assessed with a 2100 Bioanalyzer and the Prokaryote Total RNA 6000 Pico Kit (Agilent). All samples had an RNA Integrity Number (RIN) >8. RNA concentration was quantified on Qubit using the RNA Broad Range assay (Invitrogen). Ribosomal RNA was depleted and libraries were constructed using the Illumina Stranded Total RNA Prep with Ribo-Zero Plus (Illumina) using 100 ng of RNA input. IDT for Illumina UD indexes were ligated to the libraries. Library quality control was performed with the Agilent DNA 1000 kit on the Bioanalyzer. The concentration of the libraries was quantified using the Qubit dsDNA high sensitivity assay. Libraries were pooled in equimolar amounts, denatured, and diluted to a final concentration of 1.5 pM. Libraries were then sequenced on the NextSeq 500 using the mid-output kit, generating 75-bp paired-end reads. Transcriptomic sequencing results represent three biological replicates with the exception of SA925 under H_2_O_2_ conditions, which represents two replicates due to one replicate failing to yield adequate reads for analysis.

#### Transcriptomic Analysis of SA1088 and SA925

Demultiplexed reads were merged within replicates and checked for quality using FastQC on forward and reverse reads separately. Reads were trimmed using the *Trim Galore!* wrapper for FastQC and CutAdapt [https://www.bioinformatics.babraham.ac.uk/projects/trim_galore/]. Trimmed reads were aligned to the complete SA925 genome using Bowtie2 and sorted using *Samtools*. Read counts per-gene were obtained with *featureCounts()* from the *Rsubread* package, using CDS features from a SA925 genome annotation file reformatted from the output of the PGAP docker image (ncbi/pgap:2021-11-29.build5742). We mapped PGAP-annotated RefSeq IDs for each CDS to UniProt to obtain gene names and GO terms. Differential expression analysis between treatments and strains were performed using DESeq2[https://github.com/mikelove/DESeq2], and plots in Figure 5 were produced using *ggplot2*[https://github.com/tidyverse/ggplot2] and *viridis* [https://github.com/sjmgarnier/viridis], with functional categories for stress response and toxicity assigned to genes via a combination of QuickGO and UniProt mappings and manual annotation based on Gaupp et al.’s 2012 review article (Gaupp et al., 2012).

## SUPPLEMENTARY MATERIALS

**Table S1. Related to Figure 1.**

*S. aureus* isolates from DFU and their mean phenotype values.

**Table S2. Related to Figure 1.**

Linear mixed models predicting each phenotype based on healing outcome.

**Table S3. Related to Figure 4.**

Flow cytometry cell counts and percentages

**Figure S1. Related to Figure 1.**

***Staphylococcus aureus* virulence phenotypes are associated with diabetic foot ulcer outcomes.** Production for each phenotype of each isolate from DFU that healed by twelve weeks (grey) or not (orange). Data represents the mean of three biological replicates each consisting of three technical replicates. Independent t-tests were performed comparing maximum isolate phenotypes from each DFU which was healed (left) and unhealed (right) at week 12. **(A)** biofilm production, **(B)** siderophore production, **(C)** staphylokinase production, and (**D)** staphyloxanthin production.

**Figure S2. Related to Figure 2.**

Growth curves of clinical and USA300 LAC isolates. Each isolate was grown in liquid culture shaking for 8 hours (x-axis) and OD600 (y-axis) was measured every hour. Dots represent the mean of three independent experiments. Error bars depict standard deviation from the mean. **Figure S3. Related to Figure 3.**

**(A)** Pilot experiment comparing wound size 14 days post wounding for mice infected with SA925 (magenta), SA1088 (teal) and PBS control (blue). Dots represent the mean of 3 area measurements for each wound. **(B)** Kaplan-Meier survival curve comparing mouse survival across groups from all three independent diabetic mouse wounding experiments. Dotted lines represent confidence intervals. **(C)** CFUs extracted from each biopsied wound for each group of male mice infected with *S. aureus* or control 21 days post wounding. **(D)** *S. aureus* CFUs extracted and grown on staphylococcus isolation agar from each biopsied wound for each group of male mice infected with *S. aureus* or control 21 days post wounding. For **A,C,D**, error bars represent the standard deviation from the mean. A two-sided Wilcoxon sum rank test was performed amongst each group.

**Figure S4. Related to Figure 4.**

Flow cytometry gating strategy to analyze immune cell recruitment, and specifically neutrophil recruitment in response to wounding and infection with *S. aureus* strains.

**Figure S5. Related to Figure 5.**

Staphyloxanthin production by WT *S. aureus* and Sigma B operon transposon mutants. Y-axis represents percent production compared to 502A strain. Error bars represent standard deviation.

