## Supplementary figures and images for "Staphyloxanthin production by *Staphylococcus aureus* promotes resistance to oxidative stress to delay diabetic wound healing"

### Supplemental Figures

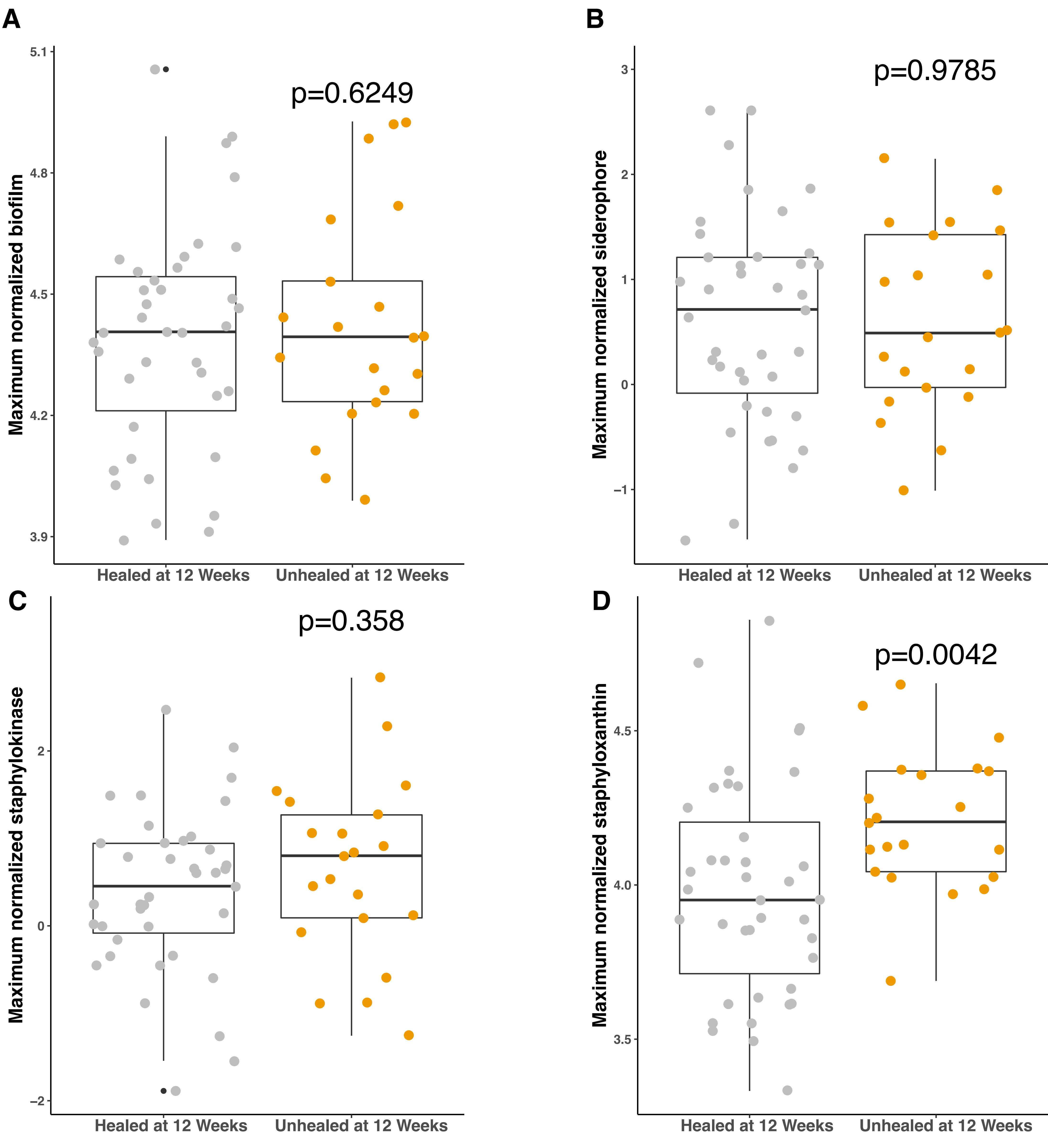

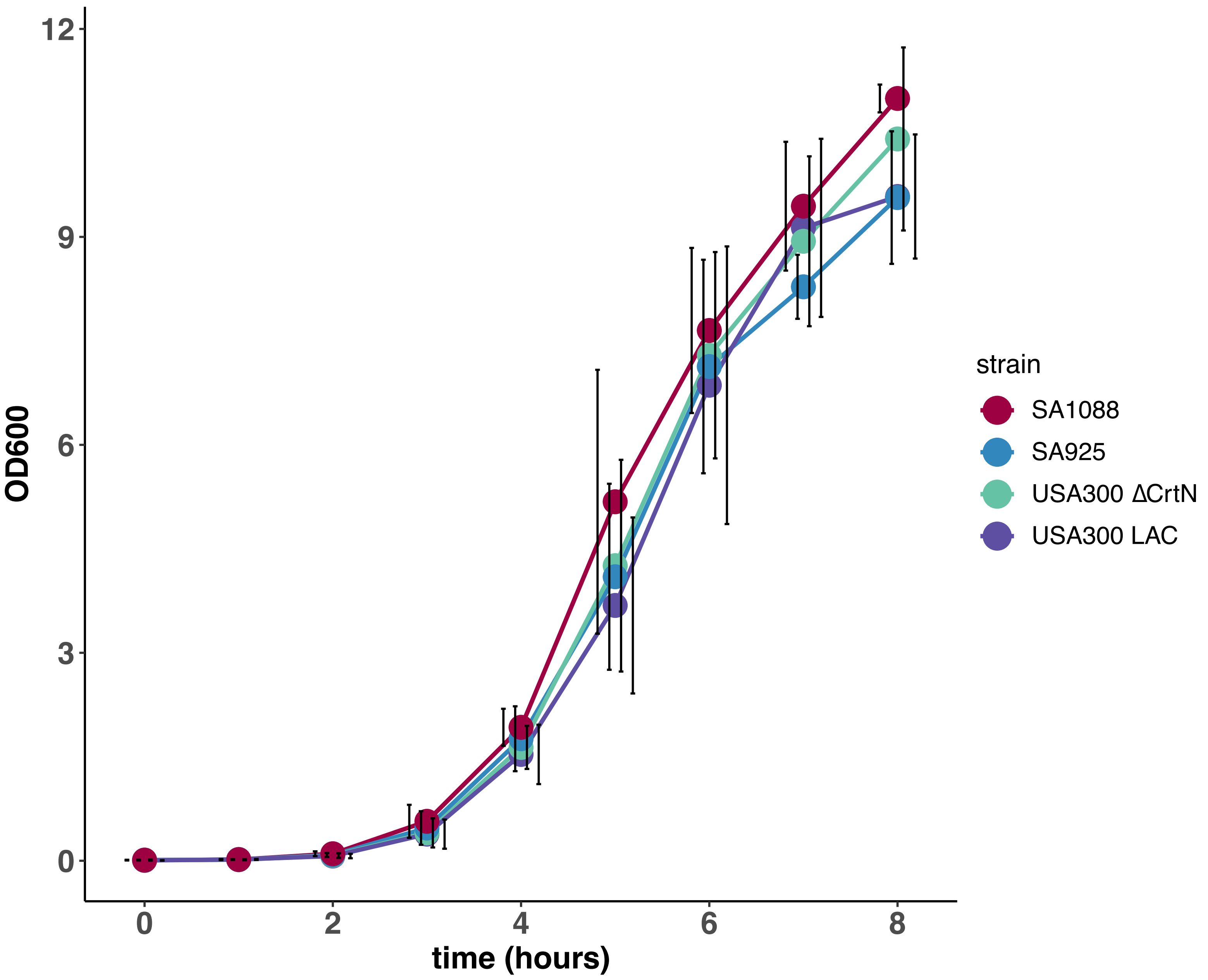

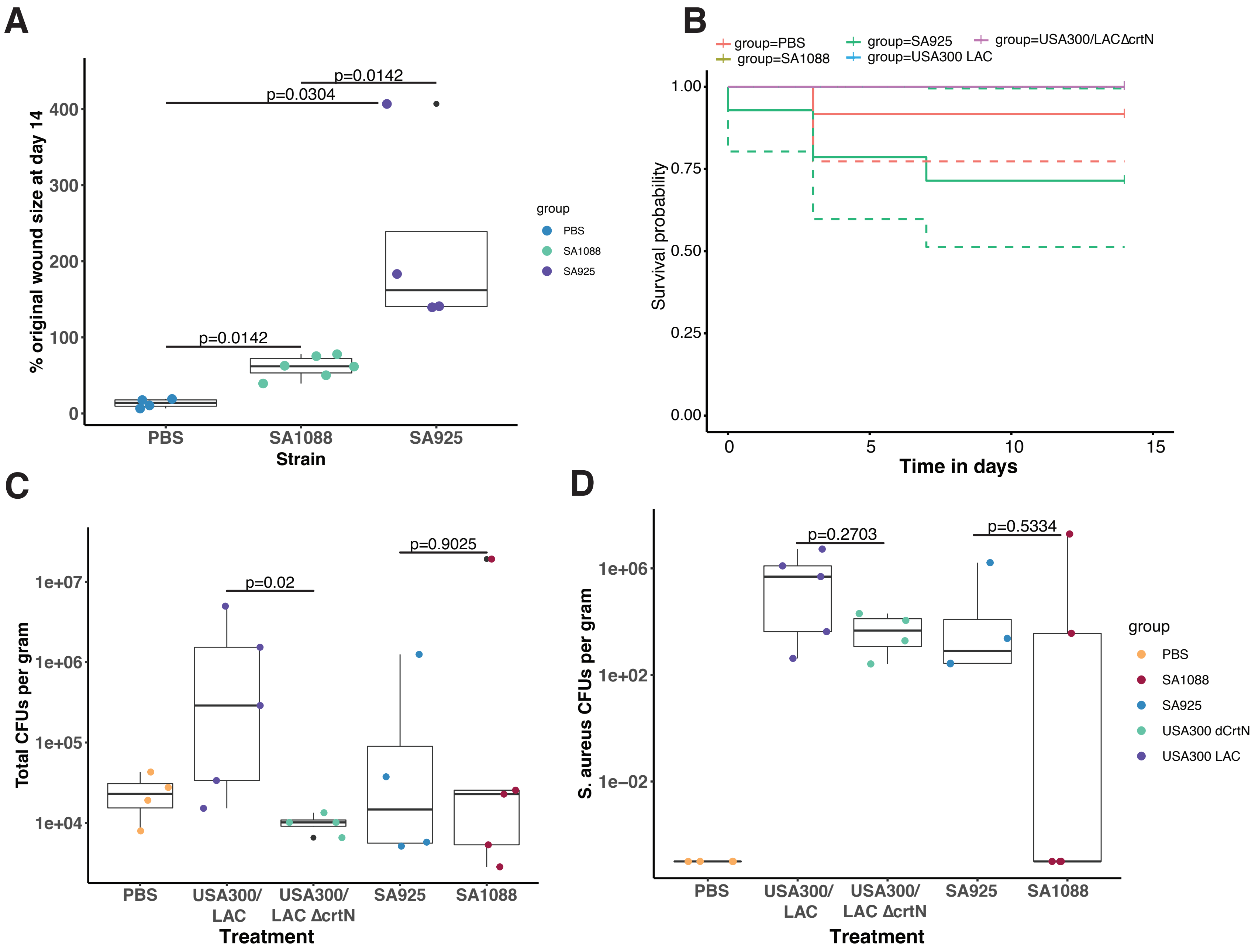

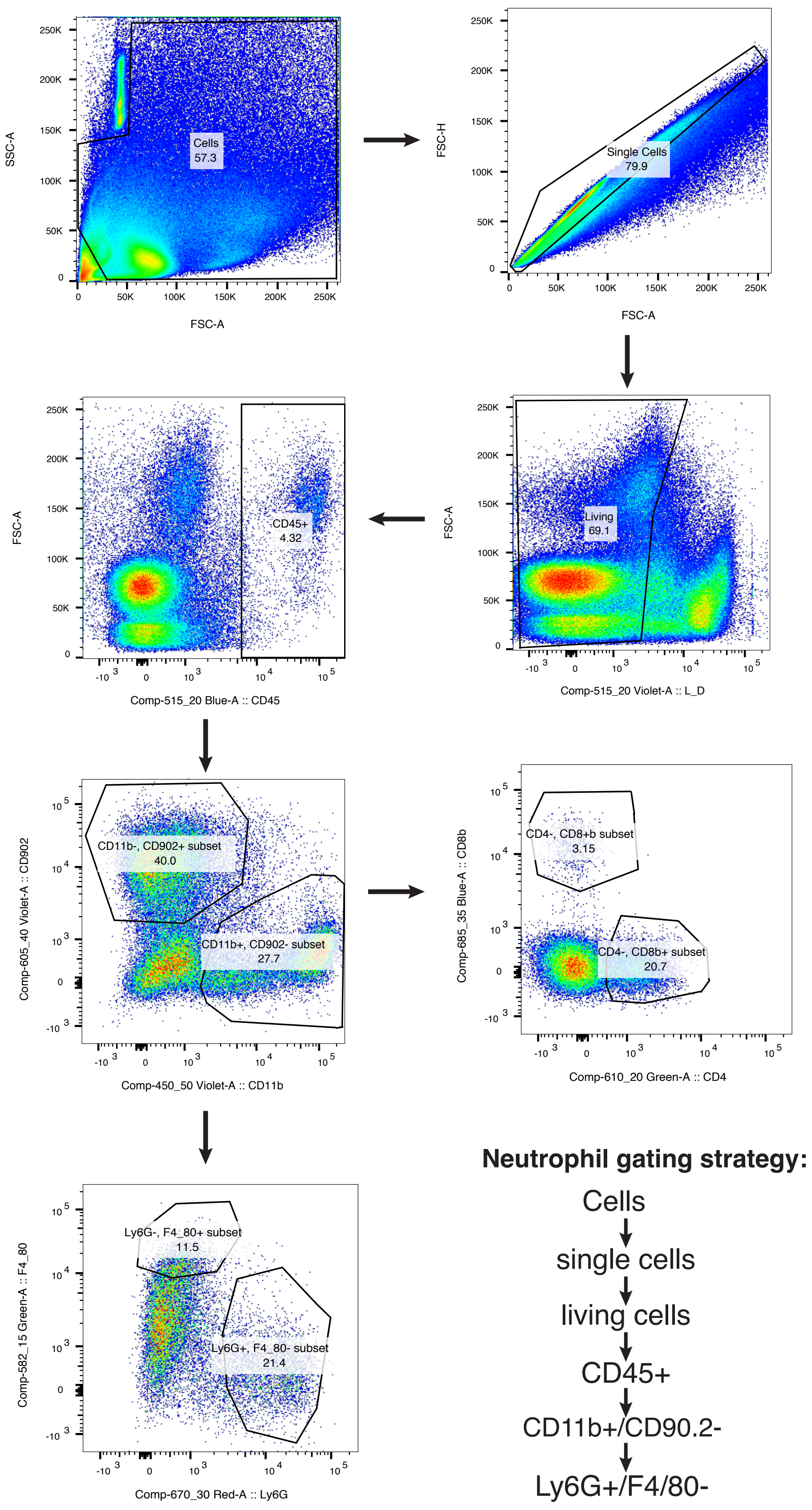

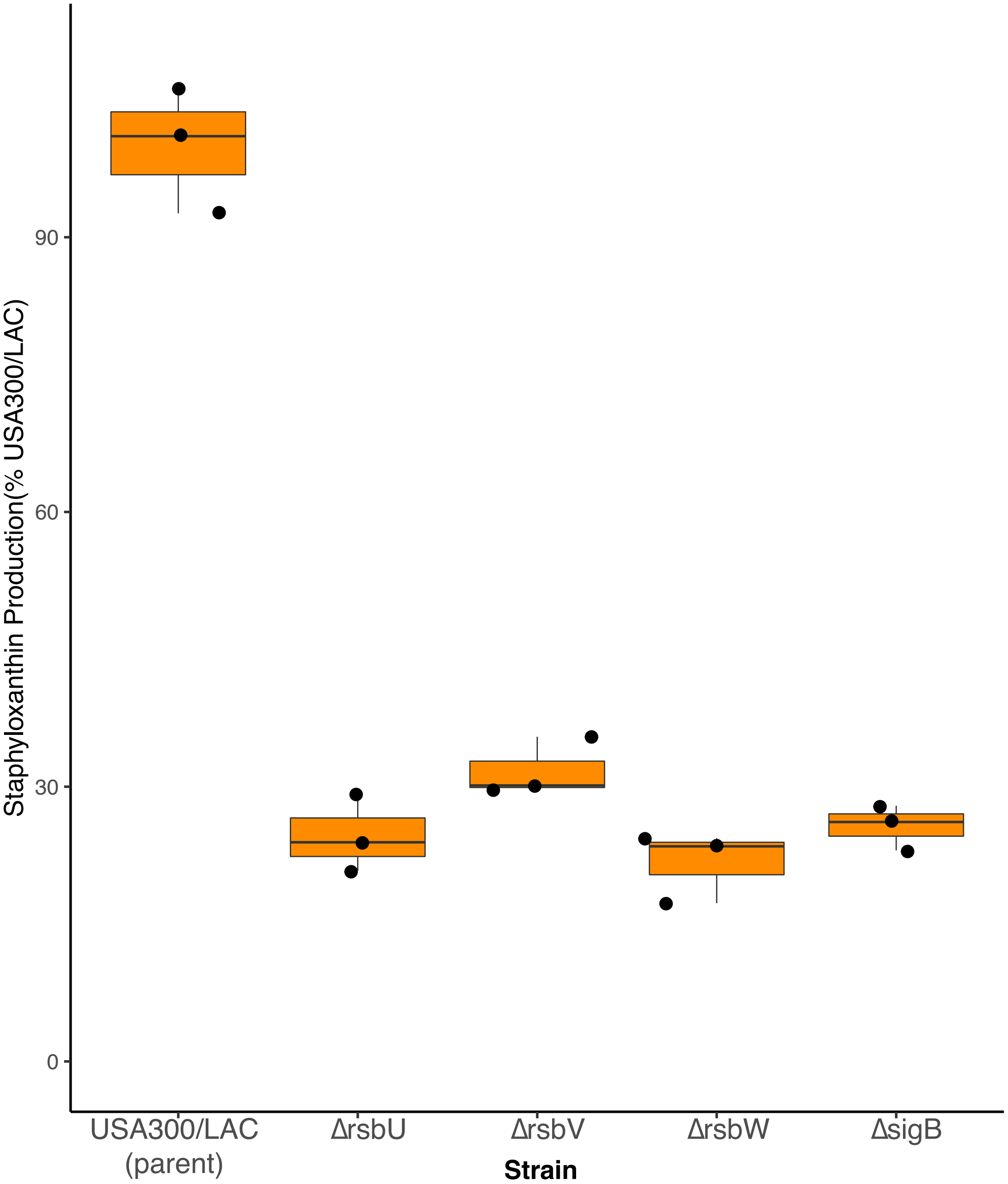
